# Familial t(1;11) translocation is associated with disruption of white matter structural integrity and oligodendrocyte-myelin dysfunction

**DOI:** 10.1101/657163

**Authors:** Navneet A. Vasistha, Mandy Johnstone, Samantha K. Barton, Steffen Mayerl, Bhuvaneish T. Selvaraj, Pippa A Thomson, Owen Dando, Ellen Grünewald, Clara Alloza, Mark E. Bastin, Matthew R. Livesey, Kyriakos Economides, Dario Magnani, Paraskevi Makedonopolou, Karen Burr, David J. Story, Douglas J. Blackwood, David J.A Wyllie, Andrew M. McIntosh, J. Kirsty Millar, Charles ffrench-Constant, Giles E. Hardingham, Stephen M. Lawrie, Siddharthan Chandran

**Affiliations:** Centre for Clinical Brain Sciences, The University of Edinburgh, Chancellor’s Building, 49 Little France Crescent, Edinburgh EH16 4SB; MRC Centre for Regenerative Medicine, The University of Edinburgh, 5 Little France Drive, Edinburgh EH16 4UU; Division of Psychiatry, The University of Edinburgh, Royal Edinburgh Hospital, Edinburgh EH10 5HF; Institute of Genetics and Molecular Medicine, The University of Edinburgh, Western General Hospital, Crewe Road, Edinburgh EH4 2XU; Centre for Discovery Brain Sciences, The University of Edinburgh, Hugh Robson Building, 15 George Square, Edinburgh EH8 9XD; Translational Sciences at Sanofi, Chilly-Mazarin, France; Centre for Brain Development and Repair, Institute for Stem Cell Biology and Regenerative Medicine, Bangalore 560065; UK-Dementia Research Institute, The University of Edinburgh, Chancellor’s Building, 49 Little France Crescent, Edinburgh EH16 4SB

**Author notes:** Biotech Research and Innovation Centre, Ole Maaløes Vej 5, Copenhagen N 2200, Denmark (Navneet A. Vasistha). Correspondence: Prof. Siddharthan Chandran, Centre for Clinical Brain Sciences, Chancellor’s Building, 49 Little France Crescent, Edinburgh EH164SB.

**Keywords:** iPS, DISC1, myelin, OPC, schizophrenia

## Abstract

Although the underlying neurobiology of major mental illness (MMI) remains unknown, emerging evidence implicates a role for oligodendrocyte-myelin abnormalities. Here, we took advantage of a large family carrying a balanced t(1;11) translocation, which substantially increases risk of MMI, to undertake both diffusion tensor imaging (DTI) and cellular studies to evaluate the consequences of the t(1;11) translocation on white matter structural integrity and oligodendrocyte-myelin biology. This translocation disrupts among others the *DISC1* gene which plays a crucial role in brain development. We show that translocation-carrying patients display significant disruption in white matter integrity compared to familial controls. At a cellular level, we observe dysregulation of key pathways controlling oligodendrocyte development and morphogenesis in induced pluripotent stem cell (iPSC) case derived oligodendrocytes. This is associated with reduced proliferation and a stunted morphology *in vitro*. Further, myelin internodes in a humanized mouse model that recapitulates the human translocation as well as after transplantation of t(1;11) oligodendrocyte progenitors were significantly reduced compared to controls. Thus we provide evidence that the t(1;11) translocation has biological effects at both the systems and cellular level that together suggest oligodendrocyte-myelin dysfunction.

## INTRODUCTION

Schizophrenia (SZ) and other major mental illnesses (MMI) such as bipolar and major depression show high heritability. Accumulating evidence from GWAS studies points to a multifactorial polygenic inheritance, with individual genes conferring a modest increased susceptibility, as well as pleiotropy^1^. In contrast, rare genetic variants, such as a balanced chromosomal translocation in a large Scottish family, that co-segregate with MMI show highly penetrance^2–6^. The range of psychiatric phenotypes observed in people carrying the balanced t(1:11) translocation suggests its study will be of considerable value for improved understanding of biological processes underlying MMI. This translocation disrupts the *DISC1* gene and segregates with schizophrenia and affective disorders in this large family

Despite multiple lines of evidence from pathological, gene expression and radiological studies, the role of glia is understudied^2,7–15^. As oligodendrocytes enable rapid impulse propagation and provide trophic and metabolic support to axons^16–19^, their dysfunction is likely to result in altered neuronal homeostasis. Further, the sole protein-coding gene disrupted by the t(1;11) translocation; *Disc1*, is known to affect specification and differentiation of oligodendrocytes^20–22^ in animal models. Hence a direct study of the impact of the t(1;11) translocation on oligodendrocytes is important. Furthermore, given the limitations of animal models of neuropsychiatric disorders, complementary models of human glia in the context of MMI are particularly necessary. Indeed, recent studies using patient derived iPS cells have shown impaired glial maturation suggesting a causal link with schizophrenia^23, 24^.

A powerful approach to interrogate the structural and cellular white matter consequences of the t(1;11) translocation is to undertake combined water diffusion MRI (dMRI) and biological studies of iPSC-derived oligodendrocytes from patients carrying the t(1;11) translocation. Previous studies using iPSCs from individuals with *DISC1* mutations and MMI have predominantly studied neural precursor and/or neuronal processes^2,25,26^. Here, we demonstrate abnormalities of white matter integrity using brain dMRI in t(1;11) translocation carrying cases compared to familial controls. In addition, case iPSC-derived oligodendrocytes display cellular and structural abnormalities *in vitro* as well as upon transplantation into hypomyelinated mice. This study elucidates a cell biological basis of oligodendrocyte-myelin deficits in MMI. In addition, we establish a human platform for future mechanistic studies.

## RESULTS

### Global changes in white matter structure and connectivity due to the t(1;11) translocation

To begin to understand the effect of t(1;11) translocation at a whole brain as well as cellular level, we used a multi-tiered approach integrating patient and control whole brain imaging, and *in vitro* and *in vivo* stem cells along with transgenic studies (Figure 1a).

**Figure 1:**
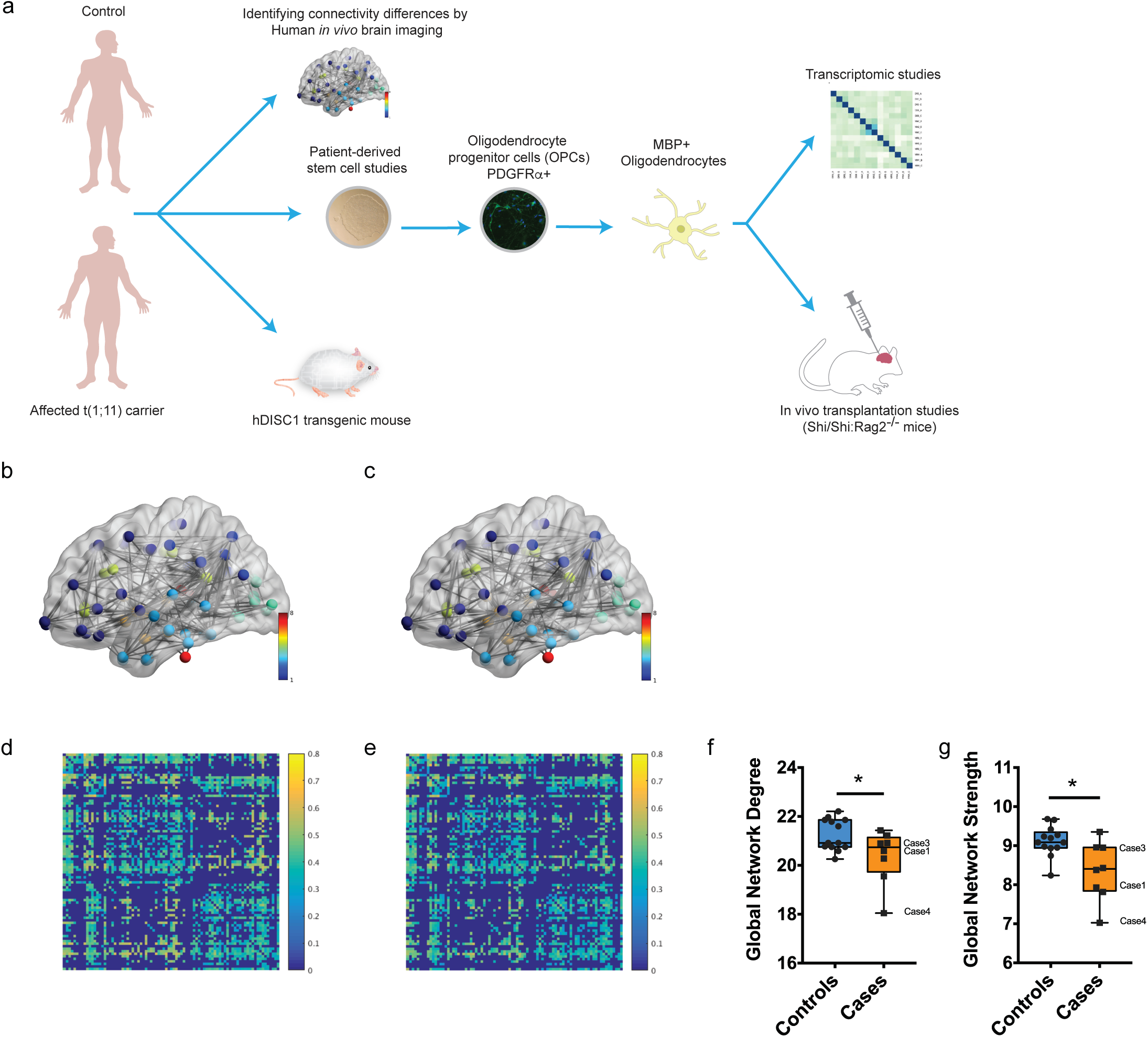
Global changes in structural networks and connectivity in t(1;11) carriers. (a) Overview of the approach used in this study to investigate the effect of t(1,11) on oligodendrocyte development and myelination capability. Brain imaging of control and affected t(1,11) carriers was undertaken to identify differences in connectivity. Separately, patient derived iPSCs were differentiated towards a oligodendroglial fate to study developmental differences. A combinatorial approach of cellular phenotyping along with transcriptomic analysis was employed. Finally, the myelination capability was studied in transgenic mice (*Der1/+*) recapitulating the human mutation as well as by transplantating hiPS derived OPCs into a hypomyelinated mouse model (*MBP^Shi/Shi^;Rag2^-/-^*). (b-e) Average FA-weighted structural networks (a, b) and connectivity matrices (c, d) for controls (b, d) and carriers (c, e) obtained using diffusion and structural MRI. Network nodes were identified from high-resolution T_1_-weighted volume scans using Freesurfer (http://freesurfer.net), with connecting edges created using probabilistic tractography (bedpostx/probtrackx; http://fsl.fmrib.ox.ac.uk/fsl) from high angular resolution diffusion MRI data. In (b, c) network nodes are coloured to indicate grey matter in different lobar structures, while in (d, e) stronger connections between different cortical regions are indicated by lighter yellow colours. Careful examination of the connectivity matrices show that a number of connections have higher FA in (d) than (e) suggesting that brain connectivity is increased in controls compared with carriers. (f,g) Quantification of Global network degree (f) and Global network strength (g) from (d,e) shows statistically significant changes in carriers (n= 13 controls and 8 translocation carriers, Markov chain Monte Carlo (MCMC) test). Median with upper and lower quartiles is shown. The whiskers depict the range. Cases studied using iPS cells in this particular study have been highlighted

To study the impact of t(1;11) translocation on global white matter structure and structural connectivity we undertook whole-brain probabilistic tractography using dMRI on 21 individuals from the previously reported Scottish family known to carry the t(1:11) translocation of whom 8 were carriers of the t(1;11) translocation^27^. All 8 carriers had a psychiatric diagnosis; 1 with schizophrenia, 4 with major depression(MDD), 3 with cyclothymia while only 1 of the 13 family members who did not have the t(1;11) translocation, had a psychiatric diagnosis^27^. The affected non-carrier however is described to carry modifier loci on chr11q2 and chr5q that might contribute to the development of MMI (described in^28^). High resolution T1-weighted structural and dMRI data were combined to create structural connectivity matrices of each participant’s brain. In these matrices (Figure 1b-e), the 85 grey matter regions, parcellated from the structural MRI data, are the nodes of the brain structural network while the connecting white matter pathways, identified from whole brain dMRI tractography, are its edges. The edge connection strength between nodes is obtained by recording the mean fractional anisotropy (FA), a measure of white matter microstructure, along tractography streamlines connecting all ROI (network node) pairs (Figure 1b-e). Global graph theory measures of brain structural connectivity, such as network degree (number of connections one node has to other nodes), network strength (the average sum of edge connections per node) and global efficiency (the average of the inverse shortest path length between nodes), can then be calculated for each subject and compared across populations to assess differences in brain structure. In our case, we hypothesised that carriers have reduced connectivity (e.g. lower degree, strength and global efficiency) than non-carriers. We estimated the effect of translocation status (carrier versus non-carrier) and age on global connectivity measures using a Markov chain Monte Carlo (MCMC) approach^29^. Significant differences between carriers and non-carriers were found for global network degree (posterior mean=-1.81; 95% CI= [−3.43, −0.34]; pMCMC=0.02) and global network strength (posterior mean = −1.64; 95% CI= [−3.09,-0.22]; pMCMC=0.03) (Figure 1f,g). Global network efficiency showed a tendency towards significance (posterior mean=-1.19; 95% CI= [−2.65, 0.26]; pMCMC=0.09]) while clustering coefficient was not significantly different between groups (posterior mean= 0.93; 95% CI= [−2.64, 0.59]; pMCMC>0.05) (Data not shown). Together these findings show that carriers have reduced structural connectivity compared with controls and are consistent with t(1;11) translocation causing widespread changes in white matter structural integrity.

### Altered differentiation and gene expression in t(1;11) derived oligodendroglia

In order to begin to determine the cellular and molecular basis underlying the observed abnormalities in white matter integrity, we next studied oligodendrocytes from affected and unaffected individuals – see Suppl. Table 1. iPSC lines were established, using an episomal non-integrating method, from 4 individuals carrying the mutation (Case 1 - cyclothymia, Case 2 - MDD, Case 3 - MDD, Case 4 – SZ; see Supplementary table 1) and 3 unaffected family controls (Supplementary figure 1). H3K27 trimethylation staining indicated typical X-chromosome inactivation in the female iPSC lines as seen by distinct foci in the nucleus while the male lines showed diffuse staining (Supplementary figure2). We generated enriched oligodendrocyte lineage cells using a previously described protocol^30^ (Figure 2a-c). Quantitative analysis showed no difference in progenitor specification as assessed by OLIG2^+^ staining on day 1 (Figure 2d). Developmentally, OLIG2^+^ neural precursors in the cortex give rise to PDGFRα^+^ oligodendrocyte precursor cells (OPCs) which differentiate into O4^+^ and MBP^+^ pre-myelinating oligodendrocytes *in vitro.* In contrast, there was a significant reduction in PDGFRα cells in cases compared to controls at 7 days post plating (Controls: 8.27±0.50% Cases: 5.61±0.71%, n=3 independent conversions, p<0.05 unpaired t-test) and a concomitant significant increase in O4^+^ cells in case derived lines compared to controls at 3 weeks differentiation (Controls: 63.42±1.3% Cases: 73.7±2.5%, n=3 independent conversions, p<0.05 unpaired t-test) suggesting increased differentiation of OPCs in case lines (Figure 2e,f). On analyzing the case lines individually, we observed that the reduction in PDGFRα^+^ cells and increased in O4+ cells was limited to Case3 and Case4 and not seen in other lines (Supplementary Figure 3a-c). We confirmed the reduction in proliferation in Case3 and Case4 OPCs by labelling dividing cells with EdU and co-staining for PDGFRα (Supplementary Figure 3d-f). Control and Case lines showed equal proportions of TUJ1+ or GFAP+ cells at 3 weeks (Supplementary figure 4a,c). Case lines however had a lower proportion of proliferating Ki67+ cells at the end of week 3 (Controls: 12.30±2.8% Cases: 5.54±1.2%, n=3 independent conversions, p<0.05 unpaired t-test), in keeping with premature differentiation (Supplementary figure 4b,c). A short-term decrease in expression of full length DISC1 mRNA has been previously linked to premature neural precursor differentiation^26, 31^. We found that full-length DISC1 mRNA expression was significantly reduced in case iPSCs and case-derived OPCs compared to control samples (OPCs: Controls: 0.72± 0.10, Cases: 0.22± 0.05, n=3 independent conversions, p<0.05 One-way ANOVA with Holm-Sidak’s correction) (Supplementary figure 4d).

**Figure 2:**
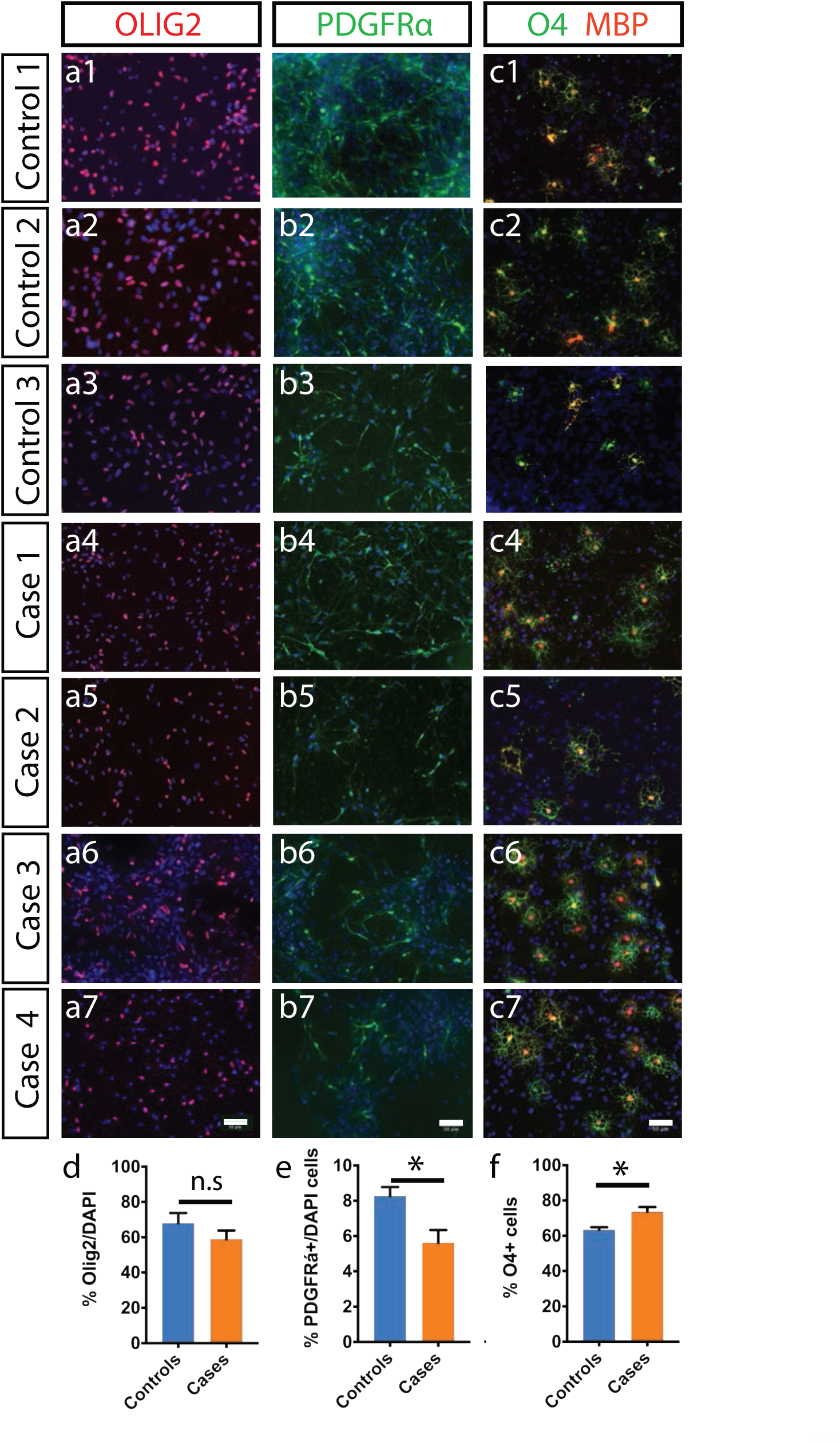
Efficient conversion of control and case iPS cells to oligodendroglial lineage. (a1-a7) Both control and case iPS lines could be patterned to OLIG2+ precursor cells under conditions described in Methods. (b1-b7) Further differentiation of cells from (a) gave PDFGRα+ OPCs under proliferative conditions containing FGF2 and PDGF-AA (c1-c7) Removal of mitogens produced mature oligodendrocytes that co-stained for O4 and MBP. (d-f) Quantification of OLIG2+ cells on day1 (d) PDFGRα+ OPCs on day7 (e) and O4+ oligodendrocytes at day21 (f) shows a significant increase in oligodendrocytes in case lines (n= 3 independent conversions for each line, unpaired t-test used to analyze p-values) Scale: (a-c) 50 μm

To begin to understand the molecular consequences of the translocation for oligodendrocytes, we next undertook RNA-seq analysis on differentiated week 3 oligodendrocyte cultures. We compared 3 case and 2 control lines by paired-end RNA sequencing on an Illumina platform. By setting an adjusted p-value of <0.05, we identified 228 genes (164 upregulated and 64 downregulated) (Figure 3a). Gene ontology (GO) analysis identified novel candidate pathways dysregulated in MMI (Figure 3b). The differentially regulated genes included those relating to nervous system development, homophilic cell adhesion via plasma membrane, galactosylceramide biosynthetic process, activation of GTPase activity, calcium ion binding, and myelination (Figure 3b,c). Genes involved in oligodendrocyte differentiation, actin binding and ion-transmembrane transport such as *NKX2-2*, *ZNF804A*, *SLC8A3, CTTNBP2* despite not being enriched in gene ontology analysis were also found to be dysregulated in t(1;11) carrying lines (*NKX2.2* Controls: 1.0±0.46 Cases: 7.16±4.41; *ZNF804A* Controls: 2.13±1.27 Cases: 25.31±24.39: *SLC8A3* Controls: 1.22±0.44 Cases: 9.56±5.71; *CTTNBP2* Controls: 1.04±0.37 Cases: 5.09±3.28, n=3 independent conversions, p<0.05 F-test of variance). Beta-actin mRNA levels however did not significantly differ between cases and controls (Supplementary Table 4). In addition, genes involved in myelin formation and wrapping such as galactosylceramide biosynthetic protein *GAL3ST1* (*GAL3ST1* Controls: 2.40±1.48 Cases: 19.98±23.76; n=3 independent conversions, p<0.05 F-test of variance) were also dysregulated in t(1;11) translocation carrying lines (Figure 3d).

**Figure 3:**
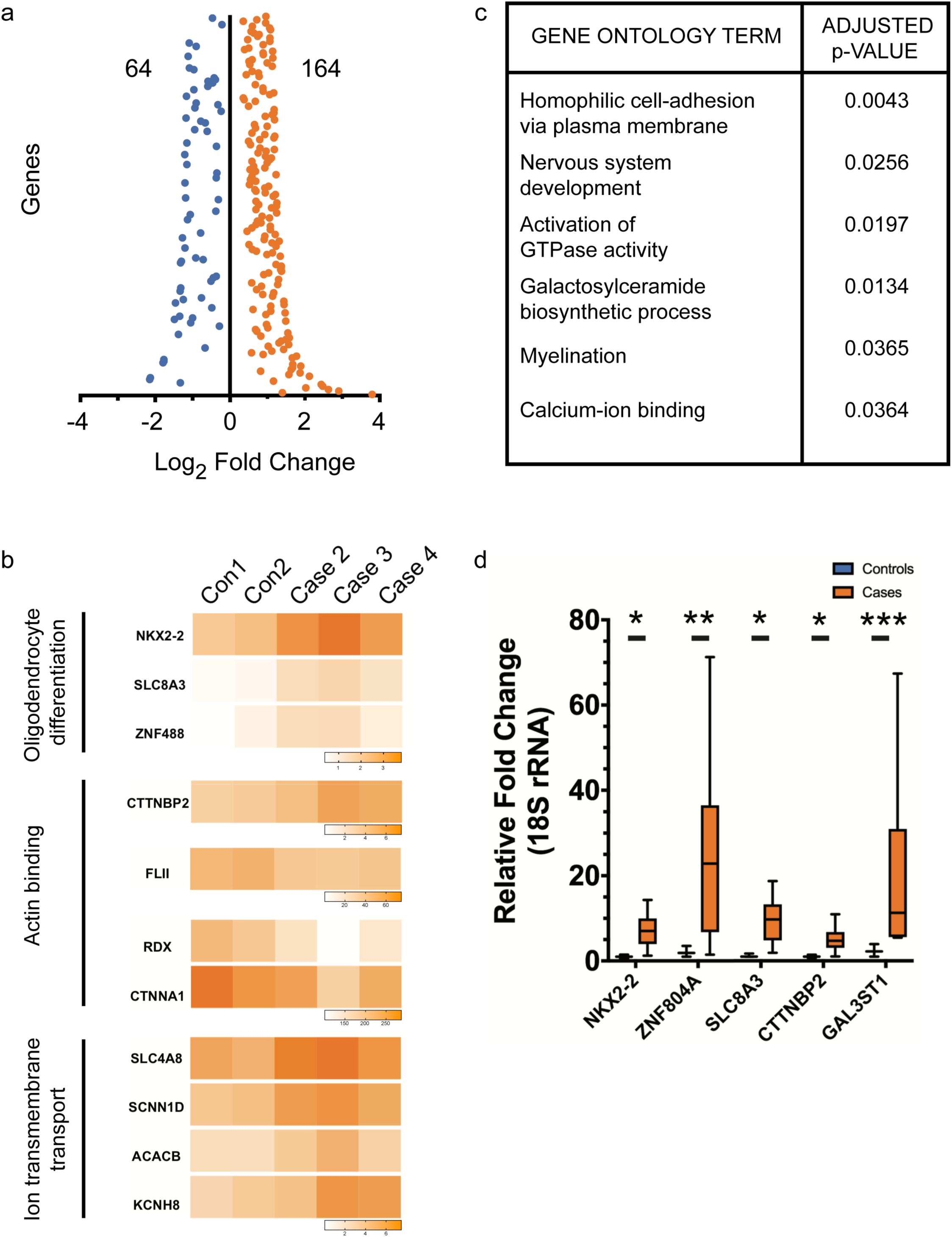
Transcriptomic analysis of Case oligodendrocytes implicate genes involved in nervous system development, activation of GTPase activity, galactosylceramide biosynthesis, myelination and calcium ion binding. (a) Differentially expressed genes (FDR 5%) were found after adjusting for multiple statistical tests. As a result, 64 and 164 genes were found down and up-regulated respectively. (b) Dysregulated pathways identified by Gene Ontology (GO) analysis of RNA-seq results. Of note are pathways implicated in nervous system development, activation of GTPase activity, galactosylceramide biosynthesis, myelination and calcium ion binding (c) Heat-maps showing differences in expression between Controls and Cases for key genes in Oligodendrocyte differentiation, Actin binding and Ion transmembrane transport. (d) qRT-PCR validation of genes selected after RNA-seq analysis (n=3 independent conversions, p<0.05 F-test for comparing variances).

### *In vitro* morphology as well as *in vivo* internodal length is severely affected in t(1;11) carrying oligodendrocytes

In support of the dysregulation of actin related genes in case lines, DISC1 has previously been shown to affect microtubule organization and neurite outgrowth^32, 33^. In oligodendrocytes, this is likely to affect morphological development and myelin formation. Case derived O4+ oligodendrocytes were found to be severely stunted and dysmorphic in comparison to familial controls (Figure 4a), a finding confirmed by quantification of surface area covered by individual oligodendrocytes (Figure 4b) (Controls: 2170μm^2^ ±179.3, Cases: 1495μm^2^ ±173.6, p<0.05 unpaired t-test, n≥30 cells per line from 3 independent conversions). The reduction in surface area was prominent across the case lines with the exception of Case1 which did not show a significant change (Supplementary Figure 5a,b). Sholl analysis also showed a severe reduction in oligodendrocyte process complexity (Figure 4c). To evaluate whether this *in vitro* phenotype might be replicated in an *in vivo* context we next examined a novel mouse model engineered to model the effects of the t(1;11) translocation upon DISC1 expression^34^. This mouse mimics the translocation-induced gene fusion that abolishes full-length *DISC1* expression from one allele and produces chimeric transcripts encoding deleterious truncated/chimeric forms of DISC1 from the derived 1 chromosome^34, 35^. Here, upon co-staining for MBP, a myelin marker, and CASPR, a paranodal marker, in mouse cortices at postnatal day 21 (Figure 4d,e), we found a significant reduction in internode lengths in the cortical layers of *Der1*/+ mutant mice (WT: 33.99 ±0.93 μm, *Der1/+*: 21.72 ±0.37 μm, n=3 mice each, >190 internodes per genotype; p<0.05 t-test with Welch’s correction) (Figure 4f). Analysis of individual myelinated axons using electron microscopy however did not show any difference in g-ratio or in the frequency of myelination (Figure 4g-i) (n=4 mice each). To assess the number of myelin sheaths produced by a single oligodendrocyte as well as their respective lengths, we performed immunofluorescence for CNPase to stain entire oligodendrocytes and hence allow for tracing processes from the cell body to myelin sheaths. Employing this technique, we analyzed oligodendrocytes in the sparsely myelinated layers II/III of the medial prefrontal cortex of 6 weeks old mice (Figure 4j). Here we found that the number of myelin sheaths produced by a single cell was significantly increased in heterozygous *Der1* mice when compared to wild-type littermates (WT: 42.57 ±2.46, *Der1/+*: 49.50 ±1.51, n=4 mice each, 28 cells per genotype; p<0.05 t-test) (Figure 4k). Additionally, in agreement with our in vitro analysis the average length of internodes produced by single oligodendrocytes was significantly reduced in heterozygous *Der1* mice (WT: 66.94 ±1.80 μm, *Der1/+*: 60.63 ±2.71 μm, n=4 mice each, 28 cells per genotype; p<0.05 t-test) (Figure 4l).

**Figure 4:**
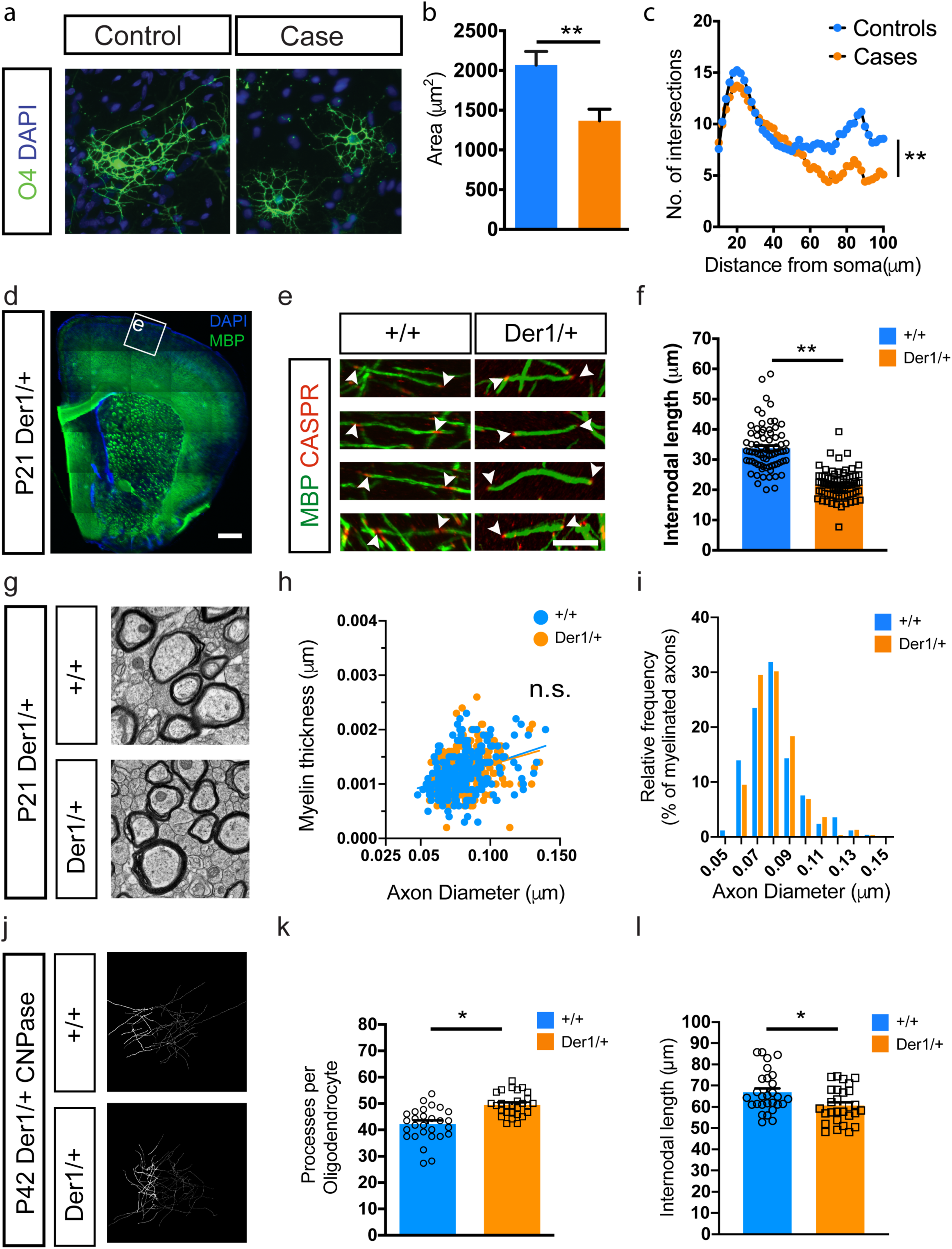
Case derived oligodendrocytes are morphologically impaired *in vitro* and, in a mouse-model recapitulating the translocation. (a) O4 staining of week 3 oligodendrocytes shows decreased cellular size and complexity of case derived cells. (b) Quantification of cellular area showing the effect of t(1;11) translocation among different case derived oligodendrocytes. n=95 for control (3 lines) and 120 for case (4 lines) oligodendrocytes. Mean ±SEM shown with paired t-test used for assessing statistical significance (p-value<0.01). (c) Case oligodendrocytes have simpler morphology with fewer branch points away from the cell soma as assessed by Sholl analysis. Mean values shown and unpaired t-test used to assess statistical significance (p-value <0.001). (d) Low magnification image of myelin basic protein (MBP) staining in the P21 *Der1*/+ cortex shows absence of gross cortical abnormalities. (Ctx: cerebral cortex, Str: striatum, cc: corpus callosum) (e) Reconstructions of myelinated segments of upper cortical layer neurons (from boxed region in (a) of wild-type (+/+) and *Der1* heterozygous (*Der1*/+) brains stained for MBP and paranodal protein Caspr. Arrowheads demarcate myelin segments used for quantification. (f) Graph showing internodal length measurements from (b). Each data point represents one myelin segment and the graph contains pooled data from n ≥ 50 segments each from 3 animals per genotype. p<0.05 Unpaired t-test with Welch’s correction. (g-i) Electron microscopy analysis of the corpus callosum of wild-type and Der1/+ mice (g) shows that thickness of myelin (h) and the relative frequency of myelinated fibers (i) are not affected due to the translocation. >250 axons measured in each group from 4 animals per genotype. (j-l) Single-oligodendrocyte analysis using by CNPase staining (j) at P42 supports findings in (e,f) and shows an increase in number of processes per oligodendrocytes (k) but a decrease in internodal length (l). Each data points represents one cell and the graph contains pooled data from n=28 cells each from 4 animals per genotype. p<0.05 Unpaired t-test. Scale bars: (a) 20μm, (c) 500μm, (d) 10μm

To next evaluate the *in vivo* consequences of t(1;11) mutation on human oligodendrocytes we established human glial chimeras by transplantation of t(1;11) and control glial cells into neonatal hypomyelinated immune-deficient mice (*MBP^shi/shi^; Rag2^-/-^*). Neonatal animals (P1-P2) were unilaterally injected with case (Case 2 and Case 4) or control (Control 1 and Control 2) derived OPCs (200,000 cells per injection) into the developing neocortex with analysis at 8- and 13-weeks post transplantation- the latest time-point allowed by animal regulations - by staining for MBP, GFAP and human nuclei (hNu). Human cells were found in the corpus callosum around the site of injection where they expressed either MBP or GFAP (Figure 5a,b). In addition, cells had also crossed the midline and differentiated. Myelin sheath lengths were greatly reduced in case derived oligodendrocytes (Figure 5c-f, Supplementary Figure 5c,d) with a majority of myelin sheaths (56% for Case2 and 48% for Case4) between 5-10μm. In comparison, only 10% of control oligodendrocytes fell in that range (Supplementary Figure 5e). Indeed, mean sheath length showed a significant difference both at 8- and 13-weeks post transplantation (8 weeks-Controls: 23.20±0.84 μm Cases: 9.48±0.26 μm, n=6 mice each and 281, 320 segments measured respectively p<0.001 unpaired t-test with Welch’s correction; 13 weeks- Controls: 29.44±1.39 μm Cases: 20.24±0.76 μm, n=3-6 mice and 269, 145 segments measured respectively, p<0.05 unpaired t-test with Welch’s correction,) (Figure 5g-h). Together the mutant mice and human transplantation studies reveal reduced myelin internode formation associated with the t(1;11) translocation.

**Figure 5:**
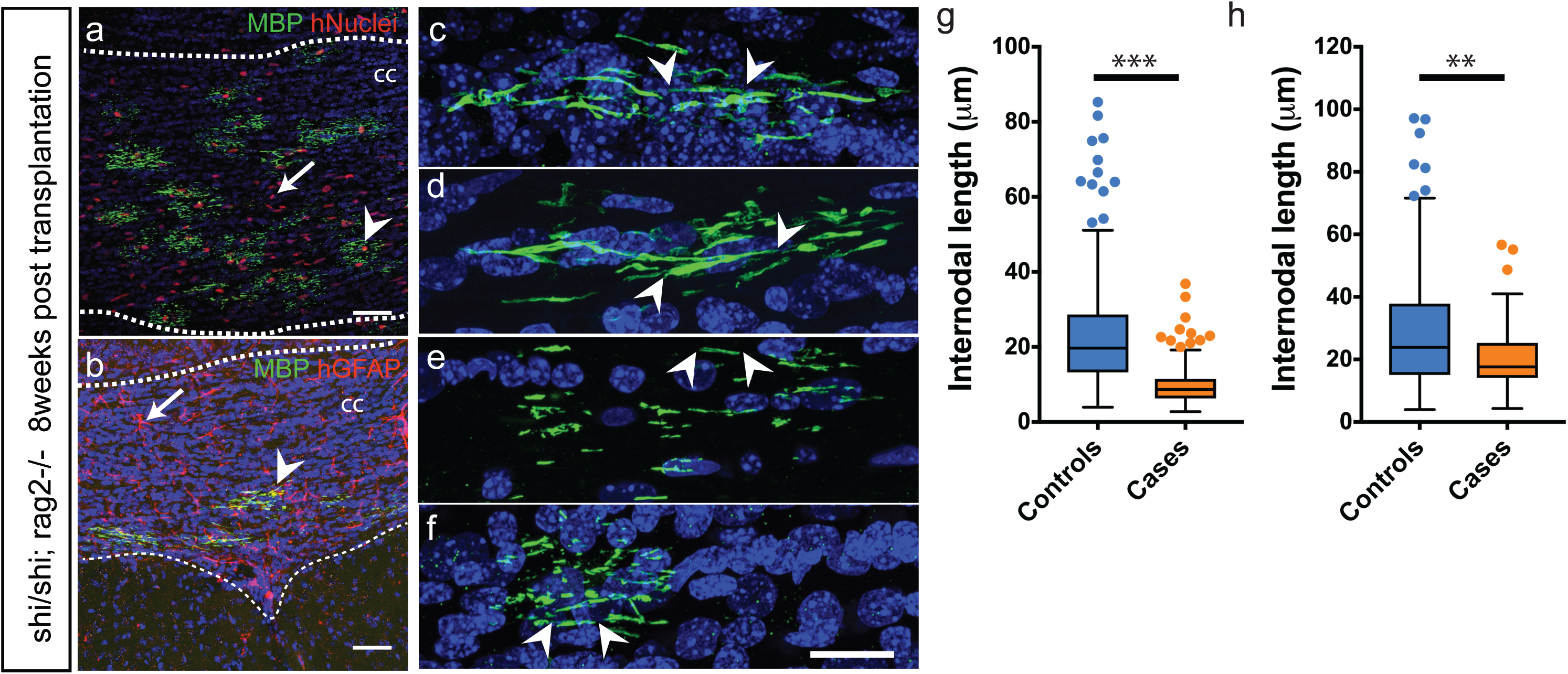
Case and control OPCs transplanted neonatally in the hypomyelinated show reduction in myelin sheath length at 8- and 13-weeks post-transplantation. (a,b) Human OPCs transplanted into *MBP^Shi/Shi^;Rag2^-/-^* neonates (at P1-P2) efficiently engraft and differentiate to oligodendrocytes (arrowhead in a) and astrocytes (arrow in b) in the white matter (dotted lines) by 8 weeks. Human cells are identified by immunoreactivity against human nuclei (red) and MBP (green) in (a) and with human GFAP (red) and MBP (green) in (b). (c-f) Individual human oligodendrocytes from Control1 and 2 (c,d) and Case2, 4 lines (e.f) were imaged and length of myelin segments measured. Arrowheads demarcate the beginning and end of individual myelin segments visualized in 3D. (g-h) Quantification of average myelin sheath lengths from (c-f) shows a severe reduction in cases at 8 weeks (g) and 13 weeks post transplantation (h). Mean with upper and lower quartiles shown. Scatter points represent outliers. p<0.05 Unpaired t-test with Welch’s correction, 8 weeks: n=3 mice injected with each cell-line were analyzed; >40 segments measured per mouse and cumulatively 281 and 320 segments measured, 13 weeks: n=1,2,3,3 mice injected with control1, control2, case2, case4 respectively were analyzed; >20 segments measured per mouse and cumulatively 269, 145 segments were measured in control and case injected mice respectively. cc: corpus callosum; hNuclei: Human nuclei; MBP: Myelin basic protein; CASPR: Contactin associated protein 2; hGFAP: Human glial fibrillary acid protein Scale bars: (a,b): 50μm, (c-f): 20μm respectively.

## DISCUSSION

In this study we provide converging lines of evidence from human imaging, molecular and cellular analyzes including of chimeric mice that suggests that the t(1;11) translocation causes oligodendrocyte-myelin deficits.

The results from structural connectivity, measured using graph theory analysis are consistent with widespread t(1;11) translocation dependent widespread changes to white matter connectivity. It has previously been shown that family members who carry the balanced translocation have a pattern of cortical thinning similar to that observed in patients with schizophrenia^36^. Here, we provide evidence disruption of white matter topology and organisation building on earlier voxel-based findings from ourselves showing a link between FA values and psychotic symptoms in those with the t(1;11) translocation^37^ and between white matter organization, genetic risk markers and cognition in the patients with ‘idiopathic’ schizophrenia scanned as part of the same project^38^. The recent large ENIGMA schizophrenia diffusion tensor imaging study of over 4000 individuals has recently shown white matter microstructural differences in schizophrenia^39^ and it is likely that the t(1;11) translocation is causing an altered regulation of schizophrenia candidate genes and the DISC1 core interactome as alternative pathways for risk in major mental illness^40^.

There has been much debate about the importance of *DISC1* as a genetic risk factor for schizophrenia^41, 42^ and it is more appropriate to instead classify *DISC1* as a genetic risk factor for major mental illness with a broad clinical phenotype. As such *DISC1* interacts with many protein partners and mechanistically affects several neurodevelopmental pathways despite not being identified as having a clear association with schizophrenia from genome-wide studies^1^. This is exemplified from the clinical phenotypes of the 4 affected carriers in this study, one with schizophrenia and three with affective disorders. Interestingly, OPCs and oligodendrocytes derived from a juvenile-onset, treatment resistant individual showed severe developmental, structural and functional changes across tests (Case 4; described in Supplementary Table 1). In contrast, cells derived from individuals presenting a less severe symptomatology (as in Case 1) also showed comparatively milder changes *in vitro*. This both highlights the need for multiple patient lines as well as the promise of iPS-based systems to model aspects of disease variation. The use of several iPS lines allows stratification based on cellular or gene expression changes as well as identification of common phenotypes to aid drug-screening approaches. In line with this, we uncovered shared gene expression changes across patient lines.

Robust and comparable differentiation of case lines compared to familial controls showed that the translocation does not interfere with early specification and patterning of cells to the oligodendroglial lineage. Case lines however, showed premature cell cycle exit of OPCs, a finding consistent with reports for cortical neurons derived from *DISC1* exon 8 interrupted iPSC lines^26^ and supported also by the observed increased expression of *NKX2.2*, *ZNF488* and *SLC8A3* genes. Of these, *NKX2.2 and ZNF488* are described to be specific to the oligodendroglial lineage as well as having a pro-differentiation effect on OPCs^43–45^. Thus, their increased expression in our dataset is consistent with the observed phenotype of precocious differentiation. Our analysis also highlights genes identified via GWAS studies and implicated in psychosis (*ZNF804A*)^46^ and suicidal behaviour (*SKA2*)^47^. Additionally, several genes (*UGT8, GAL3ST1, ZNF488, CTTNBP2, SLC8A3, GRIA4,* and *KCND2*) identified in our analysis agree with a recent study of glial progenitors from childhood onset schizophrenia patient derived iPSC lines^23^. This study of humanised glial chimeric mice suggests a potential causal role for impaired glial progenitor cell differentiation and found premature migration of GPCs, reduced white matter expansion and hypomyelination relative to controls as well as deficits in astrocyte differentiation and morphology^23^. This points towards convergent mechanisms for oligodendrocyte dysfunction in MMI despite differences in genetic basis. Future studies employing functional genetics approaches such as CRISPR and siRNA mediated knockdown could clarify the role of these genes in the pathophysiology.

Our finding of dysregulation of actin associated genes, is of considerable interest noting that *DISC1* has been shown to play a role in microtubule reorganization and neurite outgrowth^33^. In keeping with these findings, we observed case derived oligodendrocytes to have a morphological phenotype of shorter and less complex processes. Critically, we also observed in the *Der1* mouse model, that recapitulates the effects of the human translocation upon DISC1 expression, shorter internodes that was further supported by a reduction in myelin segment length found upon transplantation of case and control derived OPCs into hypomyelinated mice. Transgenic rodent models of *Disc1* have shown a range of anatomical changes including cerebral cortical thinning, reduced neurite outgrowth as well as behavioural changes (reviewed in^48^) but finer structure deficits in myelination have thus far not been shown. A crucial role for the actin remodeling pathway in myelination has been described^49–51^ and could serve as an important pharmacological target in future studies. Finally, our study also points towards a galactolipid imbalance in case derived oligodendrocytes leading to myelin defects. In keeping with this, mice genetically altered for galactolipid pathway genes present neurological manifestations due to altered axo-glia interactions and unstable myelination^52–54^. Together these findings are consistent with t(1;11) translocation dependent oligodendrocyte-myelin dysfunction.

The future evaluation of *in vivo* functional consequences of shorter internodes would be of great interest since shorter internodes are associated with reduced conduction velocities and even degenerative changes. This is particularly relevant when considering that gamma oscillations between different brain regions are known to be affected in MMI. Parvalbumin expressing interneurons, the major cell-type thought to regulate gamma oscillations are extensively myelinated^55, 56^ and its disruption could potentially contribute to the development of MMI.

In summary, our findings here of reduced white matter connectivity in t(1:11) translocation carriers add to the growing evidence of white matter deficits observed in major mental illness^57^ and suggest that oligodendroglia play an important role in the underlying pathophysiology of major mental illness.

## ACKNOWLEDGEMENTS

We thank the individuals and their families who agreed to participate in this study. We are grateful to Dr. Amanda Boyd, Mrs. Carolyn Manson and Mr. John Agnew for help with animal care and to Mrs. Nicola Clements, Mrs. Karen Gladstone and Mrs. Rinku Rajan for generous help with tissue culture and Ms. Sophie Glen for help with preparing RNA. We thank Dr. Greg Polites and Dr. Michel Didier for providing access to the Der1 mice. We are also grateful to Roslin Cells (UK) for excellent support with reprogramming of fibroblasts.

This work was supported by a Medical Research Council grant to SC, JKM and AMM and an EU 7^th^ Framework Programme Grant (607616FP7) to JKM.

NAV was supported by a Department of Biotechnology, Government of India fellowship. MJ is supported by a Wellcome Trust Clinical Career Development Fellowship and The Sackler Foundation. MRL is supported by Royal Society of Edinburgh/Caledonian Research Fund Personal Research Fellowship. The imaging work was originally supported by an award from the Translational Medicine Research Collaboration—a consortium made up of the Universities of Aberdeen, Dundee, Edinburgh and Glasgow, the four associated NHS Health Boards (Grampian, Tayside, Lothian and Greater Glasgow & Clyde), Scottish Enterprise and Pfizer.

## AUTHOR CONTRIBUTIONS

This study was conceived by NAV and SC. NAV performed all cell culture and majority of the animal related experiments including design, collection, analysis and interpretation of data. MJ analyzed medical case records, performed iPS characterization and qPCR data collection and analysis. SKB performed cell culture and qPCR analysis. SM performed single-oligonucleotide internodal length characterization and jointly performed EM analysis of *Der1* animals. EG reprogrammed human fibroblasts to iPS cells and performed characterization. CA, MB and SJL analyzed MRI and tractography data. KE designed the *Der1* mouse model. BTS, DM, MRL, KB and DJS helped with collection and assembly of data. JKM maintained the *Der1* mouse colony and was involved in iPS production. NAV, MJ and SC wrote the paper. All authors read and approved the manuscript. KE was a former employee at Sanofi. The rest of the authors declare no potential conflicts of interest

## MATERIAL AND METHODS

### Participants

Individuals with and without the t(1;11) translocation were recruited from a previously reported extended Scottish family. The t(1;11) translocation status of family members was originally ascertained by karyotyping and these were extended in subsequent studies. Researchers within the Division of Psychiatry have been in contact with members of the family for many years and through them other members of the family were invited to participate. The study was approved by the Multicentre Research Ethics Committee for Scotland. A detailed description of the study was given and written informed consent was obtained from all individuals prior to participation. A summary of the individual participants in this study, their translocation status, diagnosis and medication at the time of biopsy is given in Supplementary Table 1. Psychiatric diagnoses were established by consensus between two trained psychiatrists according to DSM-IV (TR) criteria. Diagnostic information was obtained by a face-to-face semi-structured interview using the Structured Clinical Interview for DSM-IV (SCID) supplemented by reviews of hospital records and collateral information from hospital psychiatrists and General Practitioners. The operational criteria (OPCRIT) symptom check-list was completed based on psychiatric case notes and interview data.

### MRI Imaging and Analysis

All imaging data were collected on a Siemens Magnetom Verio 3T MRI scanner running Syngo MR B17 software (Siemens Healthcare, Erlangen, Germany). For each subject, whole brain diffusion MRI (dMRI) data were acquired using a single-shot spin-echo echo-planar (EP) imaging sequence with diffusion-encoding gradients applied in 56 directions (b=1000 s/mm2) and six T2-weighted (b=0 s/mm2) baseline scans. Fifty-five 2.5 mm thick axial slices were acquired with a field-of-view of 240 mm and matrix 96 × 96 giving 2.5 mm isotropic voxels. In the same session, a 3D T1-weighted inversion recovery-prepared fast spoiled gradient-echo (FSPGR) volume was acquired in the coronal plane with 160 contiguous slices and 1 mm isotropic voxel resolution. Image processing and tractography is described in Supplementary Methods.

### Animal Ethics

All animal experiments were conducted in accordance with the UK Animals (Scientific Procedures) Act (1986). Shiverer mice (C3Fe.SWV-Mbpshi/J; 001428) were purchased from Jackson Laboratories. *MBP^Shi/Shi^; Rag2^-/-^* homozygous males were crossed with *MBP^Shi/+^; Rag2^-/-^* heterozygous females to obtain 50% double homozygous pups. Der1 heterozygous animals and control littermates were obtained by crossing wild-type males with heterozygous females. The date of vaginal plug was observed as E0.5 and embryos timed accordingly. All mice were maintained on a C57BL/6J background on a reversed 12-hour light cycle with ad libitum access to food and water.

Shiverer, Rag2 and Der1 mice were identified by genotyping from tail/ear-clips using the following primers (in 5’-3’ orientation):

**Mbp_F** TCC CTG GTG GCA GCT ATG AGC AGA CAC TGA

**Mbp_R** CCC CGT GGT AGG AAT ATT ACA TTA CCA GCT

**Shi_F** AGG GGA TGG GGA GTC AGA AGT GAG GAA AGA

**Shi_R** ATG TAT GTG TGT GTG TGC TTA TCT AGT GTA

**Rag2_F** GGT CAT CCT TTG CAA CAC AG

**Rag2_M** CAG CGC TCC TCC TGA TAC TC

**Rag2_R** TGC ATT CCT AGA GCG TCC TT

**Disc1_F** CCT GCA TCC ACA GAC GTG C

**Disc1_R** CAG TAG TAA GAA AAG AGA CAA CCC CC

**Disc1_M** ATA ACG GTC CTA AGG TAG CGA GC

### Mouse model of t(1;11) translocation

This mouse line is described in^34^. Briefly, to generate a complementary model of the translocation that mimics its effects upon DISC1 expression, the endogenous mouse *Disc1* locus was genetically engineered using Regeneron’s GEMM platform (VelociMouse®). The mouse *Disc1* allele in C57BL/6 x 129/Sv ES cell clones was targeted by deleting approximately 98.5kb of genomic DNA encompassing exons 9-13, thus removing the 3’ half of the gene that is translocated to chromosome 11 in human t(1;11) carriers. This was replaced with approximately 114kb of human chromosome 11 genomic DNA containing exons 4-7 of the *DISC1FP1* gene, corresponding to the region of the gene that is translocated to chromosome 1 in t(1;11) carriers. The edited endogenous mouse *Disc1* locus thus mimics the *DISC1*/*DISC1FP1* gene fusion event on the derived chromosome 1 in t(1;11) carriers.

### CNPase Single Oligodendrocyte tracing

For CNPase immunofluorescence studies, female mice were transcardially perfused with 4% PFA at 6 weeks of age. Brains were cut into 100 µm coronal sections on a vibratome (Leica). Free-floating sections containing the medial prefrontal cortex were blocked and permeabilized with 10% normal goat serum in 0.25% Triton X-100/PBS and subjected to antigen retrieval in 0.05% Tween20/10 mM tri-sodium citrate (pH 6.0) at 95 °C for 20 min. Subsequently, sections were washed in PBS, blocked as above and incubated with primary mouse anti-CNPase antibody (Atlas antibodies; 1:2000) in blocking buffer for 24 h at 4 °C. Thereafter, sections were washed in PBS, incubated with an Alexa Fluor 488-labeled secondary antibody raised in goat (Invitrogen; 1:000) for 24h at 4 °C, and Hoechst33258 (5 µg/ml) for 20 min to label nuclei. After final washes in PBS, sections were mounted on slides and z-stack images through the whole section were taken using a Leica SP8 confocal microscope. Images were analyzed blind to genotype by assigning random numbers to the animals and using the ImageJ simple neurite tracer plugin.

### Neonatal transplantation of OPCs into *MBP^Shi/Shi^;Rag2^-/-^* mice

Homozygous *MBP^Shi/Shi^;Rag2^-/-^* pups between P1-P2 were anaesthetized with isoflurane and maintained on a heat-mat for the duration of the transplantation. Dissociated case and control-derived OPCs at day 51 (day 0= iPSCs) were delivered (200,000 cells in 2μl) using a Hamilton syringe (Hamilton Company) through the μskull. Transplants were directed at the presumptive corpus callosum and surrounding cortex. Pups were transcardially perfused with 4% paraformaldehyde at 8- and 13-weeks post injection and brains collected for histology. 3-4 mice were transplanted per line.

### Electron Microscopy

For electron microscopy, male and female mice at the age of three weeks were anesthetised and intracardially perfused with 4% paraformaldehyde/ 2% glutaraldehyde in 0.1 M phosphate buffer. After overnight post-fixation, brains were cut into 1 mm thick coronal sections using a brain slicer. Sections between Bregma 1.0 and −1.0 were trimmed further to isolate the medial aspect of the corpus callosum in a 1 mm x 2 mm x 1 mm block (L x W x H). Tissue samples were fixed in perfusion buffer for 24 h and 1% OsO4 in 0.1 M phosphate buffer for 30 min. Using an automated processor (Leica), samples were dehydrated in rising concentrations of Ethanol, incubated in propylene oxide and rising concentrations of resin (TAAB laboratories) in propylene oxide before being embedded in resin and hardened for 22 h at 60 C. Ultrathin sections were produced and analysed with a TEM-1400 Plus (Joel).

### Induced pluripotent stem cell line maintenance

All iPSC were derived from human donor dermal skin fibroblasts using integration-free episomal methods^58^. iPS lines were maintained in Matrigel (BD Biosciences) coated plastic dishes in E8 medium (Life Technologies) at 37°C and 5% CO_2_.

### Karyotyping, pluripotency markers and X-chromosome inactivation staining

Standard G-banding chromosome analysis was performed to confirm chromosome number and gross genetic abnormalities over the course of this study. Pluripotency of case and control iPS lines was confirmed by staining for SOX2, OCT3/4 and TRA1-60. X-chromosome inactivation in female iPSC lines was confirmed by staining for H3K27 trimethylation marks^59^.

### OPC and Oligodendrocyte generation

Derivation of neural precursors (NPCs) was performed as described^30, 60^ (also described in Supplementary Methods). Briefly, NPCs were cultured in suspension as neurospheres in Advanced DMEM/F12 containing: 1% N-2 supplement, 1% B27 supplement, 1% Anti-anti, 1% GlutaMAX (all from Life Technologies), FGF2 20 ng/ml (PeproTech), PDGFα 20 ng/ml (PeproTech), purmorphamine 1 μM (Calbiochem), SAG 1 μM (Calbiochem), IGF-1 10 ng/ml (PeproTech), Tri-iodothyronine (T3) 30 ng/ml (Sigma) for 6-12 weeks. Following papain dissociation of neurospheres (Worthington), cells were plated on 1/100 diluted Matrigel (BD Biosciences), 20 µg/ml Fibronectin (Sigma) and 10 µg/ml Laminin (Sigma) coated coverslips or plates at a density of 20,000-30,000 cells per 0.3 cm^2^. Differentiation of oligodendrocytes was achieved by mitogen withdrawal (except for T3 and IGF-1).

### RNA Sequencing

Library prepraration, sequencing and analysis is described in detail in Supplementary methods. Sequencing reads will be made available via ArrayExpress or European Genome-Phenome Archive (EGA) prior to publication.

### Quantitative Real-Time PCR (qRT-PCR)

RNA was extracted from cell pellets using the RNeasy Mini kit (QIAGEN). 500ng of RNA was used to prepare cDNA by random hexamer extension using MMuLV reverse transcriptase (ThermoFisher) according to the manufacturer’s instructions. Gene specific primers were used in a 96 well plate format to quantify differences in cDNA levels using a BioRad CFX96 qPCR machine. ΔΔCt method was used to normalise and quantify relative fold changes in gene expression.

Primers used in this study were (in 5’-3’ orientation):

**DISC1_F** CCA GCC TTG CTT GAA GCC AAA A

**DISC1_R** TGA GGA GTC CCT CCA GCC CTT C

**GAL3ST1_F** CAA GAC CCG GAT CGC TAC TAC

**GAL3ST1_R** TGT CAT AGC CCA GGT CGA AGA

**SLC8A3_F** CTG CAC CAT TGG TCT CAA AGA

**SLC8A3_R** GCG TCT GCA TAT ACA TCC TGG A

**NKX2.2_F** GTC AGG GAC GGC AAA CCA T

**NKX2.2_R** GCG CTG TAG GCA GAA AAG G

**CTTNBP2_F** AAG AGC GTG GCA AGA ACA AG

**CTTNBP2_R** TGG GCC TCC TCT ATG ACT TTG

**ZNF804A_F** AGT GGC CCC ATG TTC AAA TCA

**ZNF804A_R** CCA CAA CAA CTC GTT GGG AAA T

**GAPDH_F** GAG TCC ACT GGC GTC TTC AC

**GAPDH_R** ATG ACG AAC ATG GGG GCA T

β**-ACTIN_F** CAC CTC CCC TGT GTG GAC TTG GG

β**-ACTIN_R** GTT ACA GGA AGT CCC TTG CCA TCC

### Immunocytochemistry

All steps were performed at ambient temperature. Cells were fixed with 4% paraformaldehyde in PBS for 5 minutes, permeabilized with 0.1% Triton X-100 containing PBS (except for PDGFRα and O4), blocked in 3% goat serum and incubated with appropriate primary and secondary antibodies (Alexa Fluor conjugated; Life Technologies). Nuclei were counterstained with DAPI (Sigma) and coverslips were mounted on slides with FluorSave (Merck). Fields were selected based upon uniform DAPI staining and imaged in three channels. Antibodies; OLIG2 (1:200, Millipore), PDGFRα (1:500, Cell Signalling), O4 (1:500, R&D Systems), MBP (1:50, Abcam), GFAP (1:500, DAKO), GFAP (1:500, Cy3 conjugated; Sigma), CASPR (1:1000, Abcam), Sox2 (1:1000, Abcam), Oct3/4 (1:250, Santa Cruz), Nanog (1:100, R&D Systems), TRA1-60 (1:100, StemGent), Human Nuclei (1:300, Millipore), Human GFAP (1:1000, Biolegend), Neurofilament-H (1:10,000, Biolegend).

Proportion of mitotic cells was carried out by labelling OPCs with 10μM ethynyl μdeoxy-uridine (EdU, Life Technologies) on day 3 for 24 hours. Media was replaced completely the next day and cells cultured for an additional 48 hours in the absence of EdU before fixation using 4% PFA (Sigma). For detection, cells were permeabilized using 0.1% saponin (Sigma) for 2 hours and detected using the EdU detection kit (Life Technologies) according to manufacturer’s instructions.

### Image acquisition and analysis

Images were acquired either using a Zeiss ObserverZ1 wide field or Zeiss LSM710 confocal microscope (Carl Zeiss Microimaging).

Images were converted to TIFF format, corrected for brightness and contrast and analyzed using ImageJ (NIH) or Adobe Photoshop (Adobe Inc.). Figures for preparation were assembled using Adobe Illustrator (Adobe Inc.)

### Cell morphometry

Cellular area was measured in ImageJ using O4 stained images^61^. Images were thresholded, segmented and individual cells outlined using the wand selection tool. Measurement of internodal lengths was performed using Simple Neurite Tracer plugin in ImageJ. Electron microscopy measurements were made using the freehand line tool in ImageJ. Areas was calculated by the in-built area measurement function.

### Statistical Analysis

Statistical analysis of the data was performed with Prism (GraphPad software, USA). All tests were performed as two-tailed. Tractography data was analyzed by Markov Chain Monte Carlo modelling^29^.

For experiments involving iPSCs, each derivation was considered as a n=1 and data was collected from a minimum of 3 derivations for each line. Normality of data distribution was analyzed by D’Agostino and Shapiro-Wilk’s tests. Normally distributed data was analyzed using Student’s t-test or by 1-way ANOVA test with Holm-Sidak’s multiple comparison correction (for more than 2 groups). Welch’s correction was applied when standard deviations of the two groups differed from each other. Quantitative PCR data was analyzed using non-parametric tests such as Mann-Whitney or Kruskal-Wallis tests (for 2 and >2 groups respectively).

