## Supplementary Table3 for "Familial t(1;11) translocation is associated with disruption of white matter structural integrity and oligodendrocyte-myelin dysfunction"

**SUPPLEMENTARY TABLE 3:** CASE DESCRIPTIONS AND CELL LINE IDs USED IN THIS STUDY

| Subject | Sex | Diagnosis | t(1:11) Translocation status | Psychotropic Medications (At time of biopsy) | AAO/AAS | Pedigree Reference (Ryan et al., 2018) | Pedigree Reference (Blackwood et al., 2001) |
|---|---|---|---|---|---|---|---|
| Control 1 | Male | Unaffected | Non-Carrier | None | NA/73 | 40 | IV 15 |
| Control 2 | Female | Unaffected | Non-Carrier | None | NA/52 | 28 | not included* |
| Control 3 | Female | Unaffected | Non-Carrier | None | NA/46 | 29 | not included* |
| Case 1 | Male | Cyclothymia | Carrier | None | 45/66 | 55 | not included* |
| Case 2 | Male | Major Depressive Disorder | Carrier | Fluoxetine | 37/75 | 24 | IV 11 |
| Case 3 | Male | Major Depressive Disorder | Carrier | Citalopram | 39/58 | 19 | IV 9 |
| Case 4 | Male | Schizophrenia | Carrier | Sodium valproate, Clozapine, Lithium Carbonate | 20/60 | 18 | IV 8 |

* Not included in the pedigree in Blackwood et al., 2001
