## Supplementary figures and images for "Familial t(1;11) translocation is associated with disruption of white matter structural integrity and oligodendrocyte-myelin dysfunction"

### Supplementary Figure1

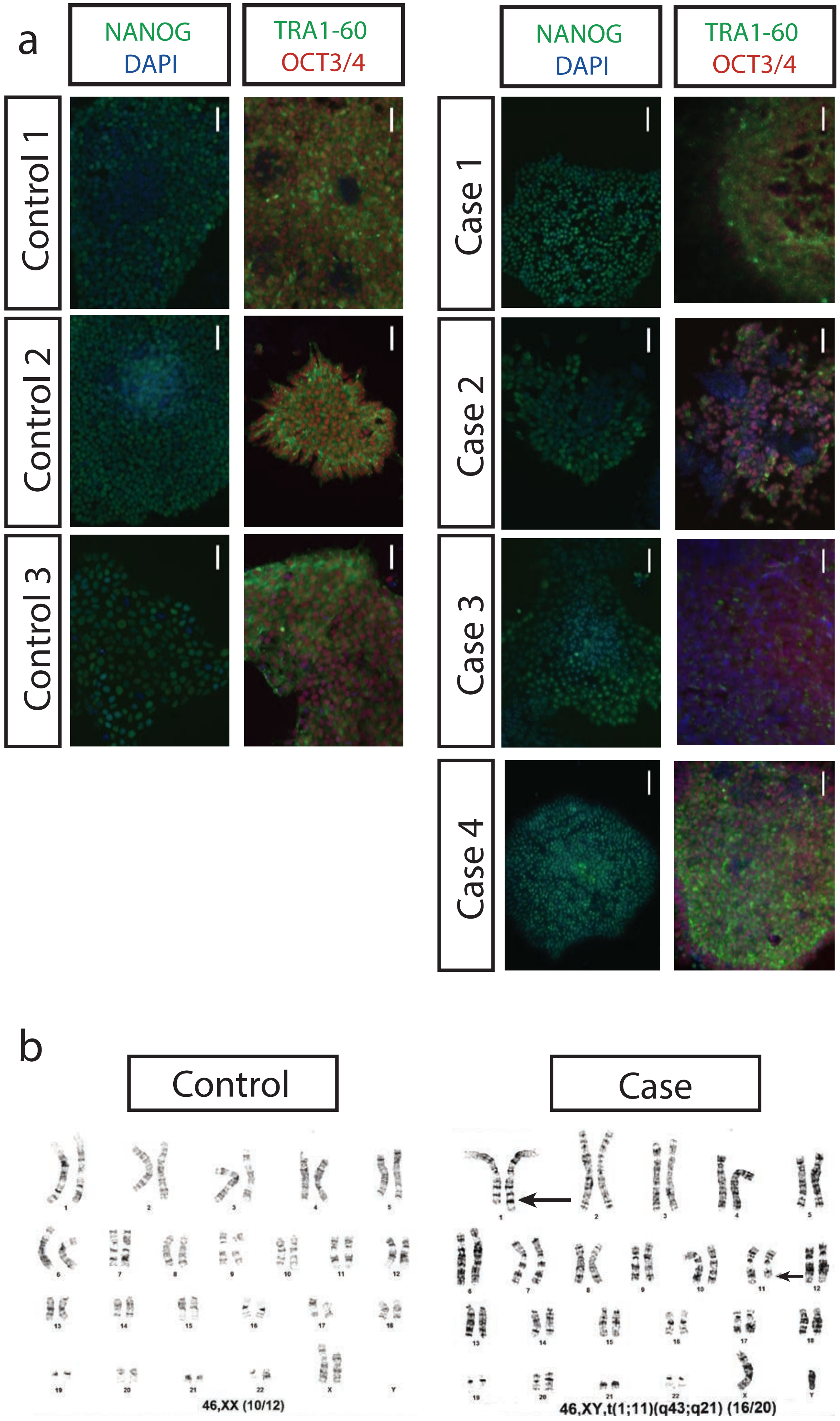

### Supplementary Figure2

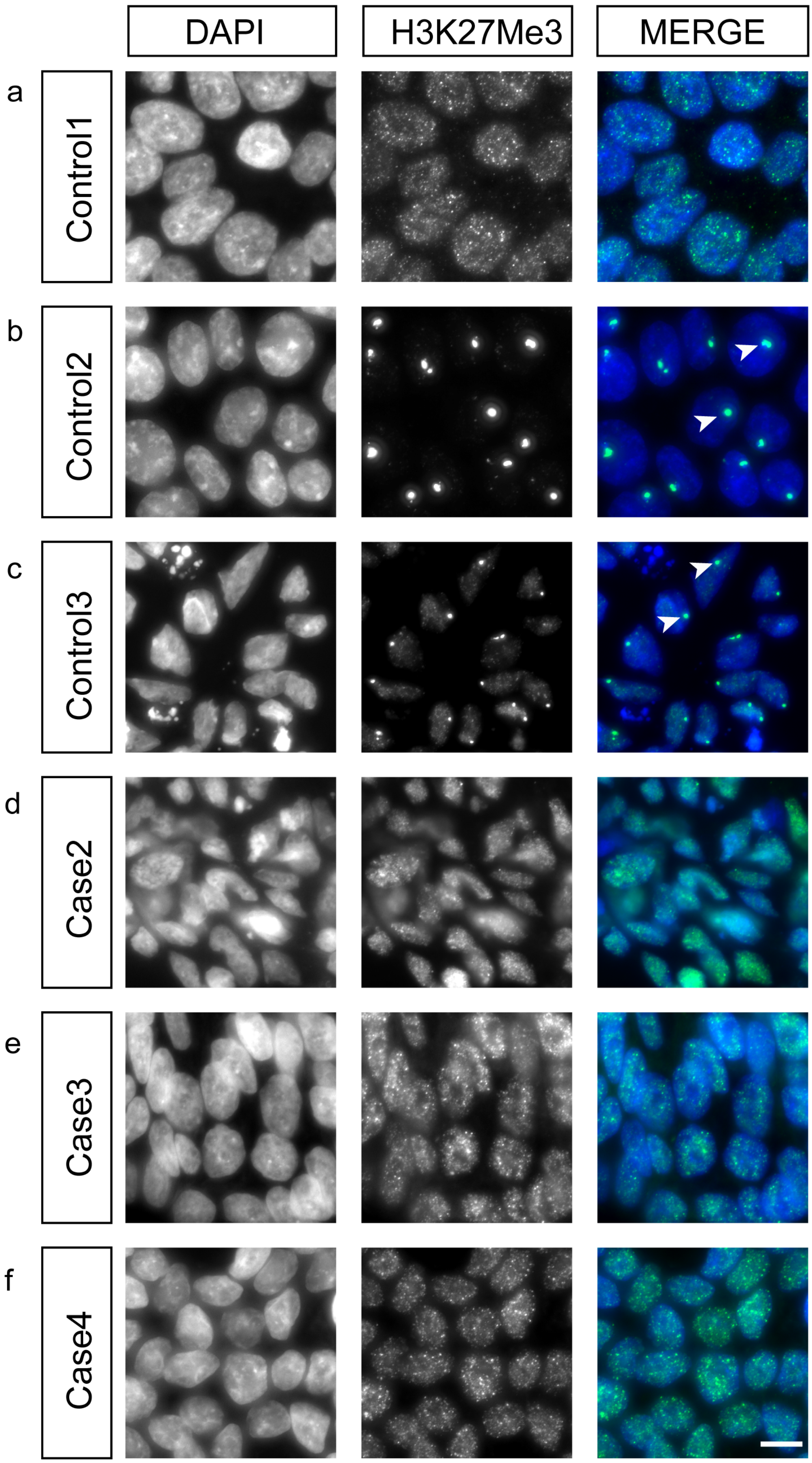

### Supplementary Figure3

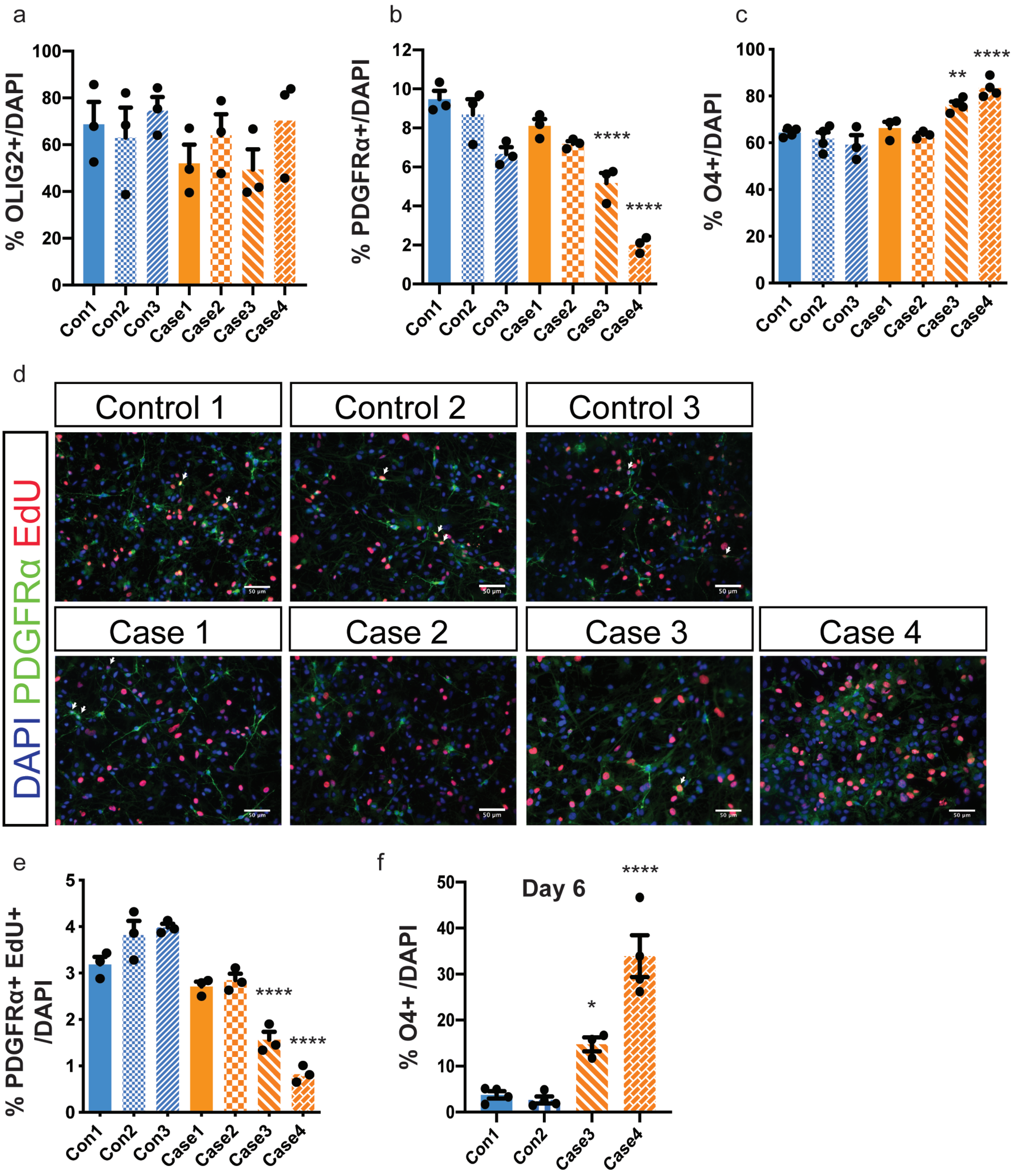

### Supplementary Figure4

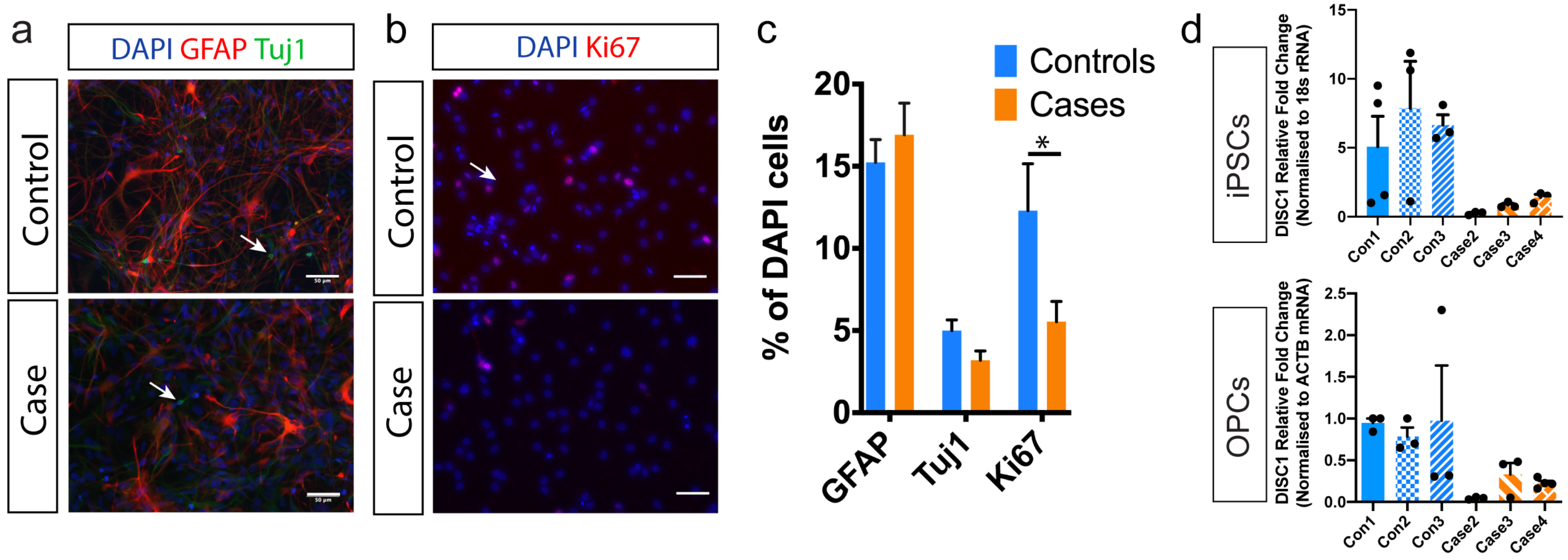

### Supplementary Figure5

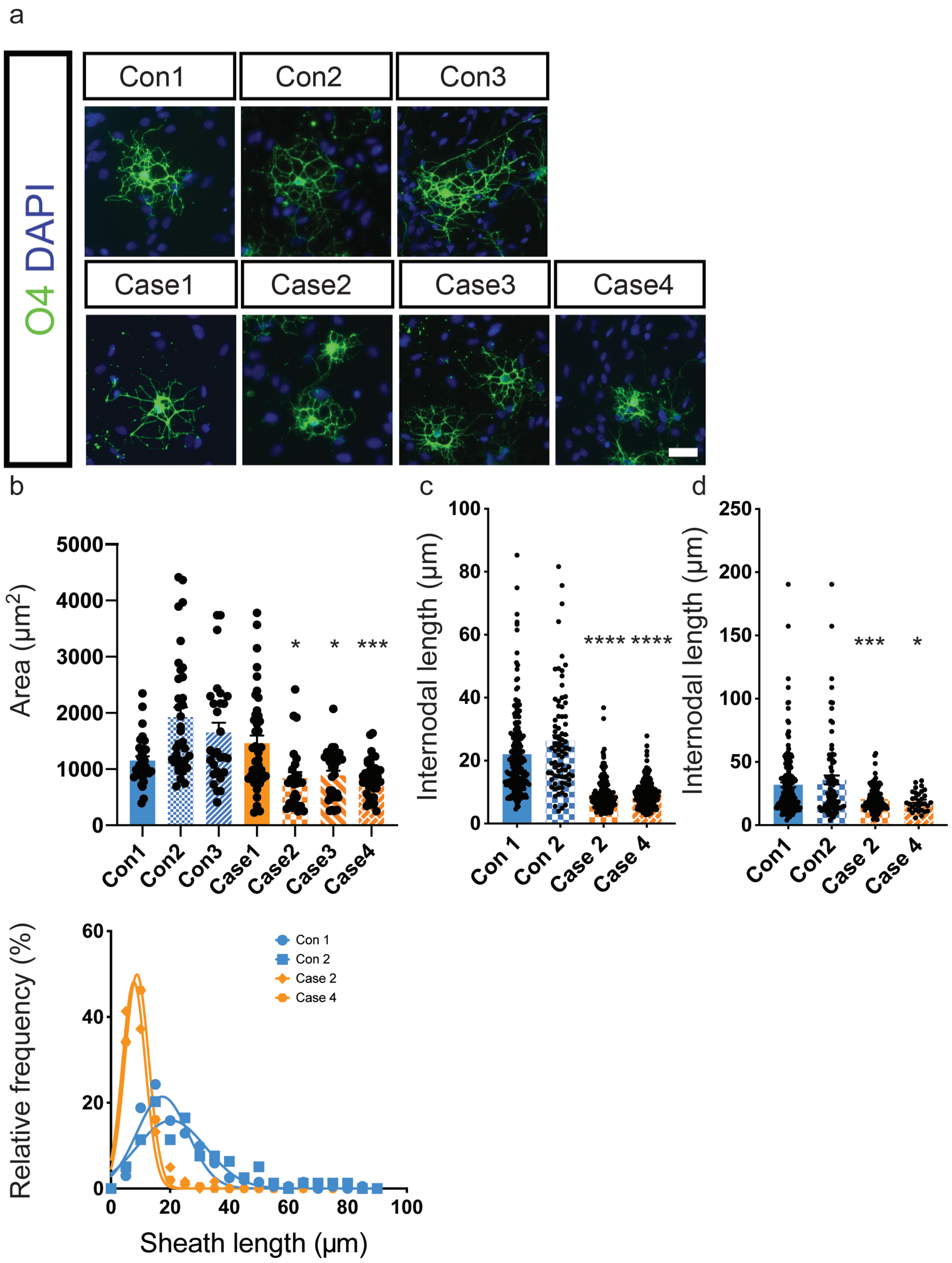

### Supplementary Figure6

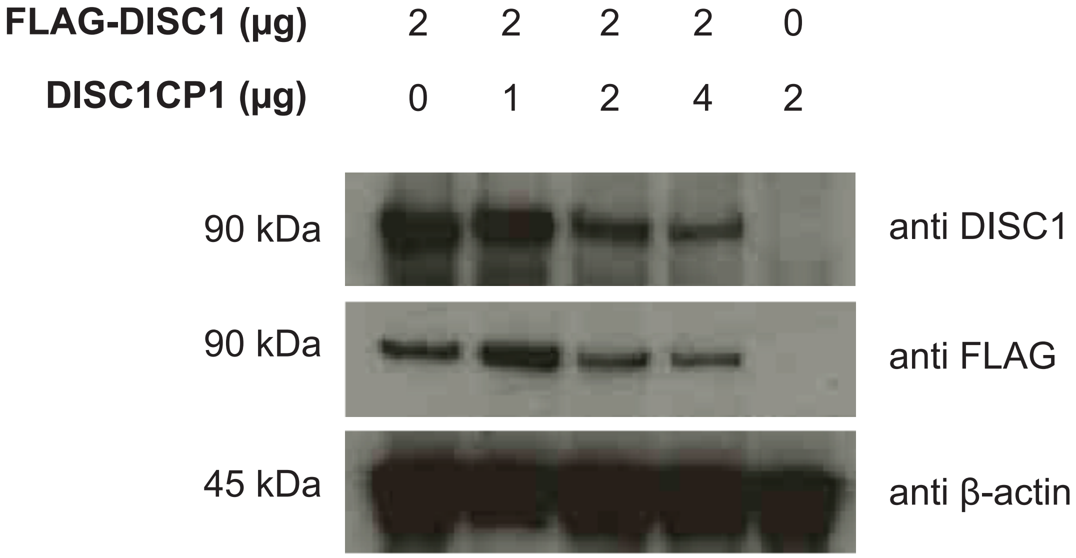
