## Supplementary Table2 for "Familial t(1;11) translocation is associated with disruption of white matter structural integrity and oligodendrocyte-myelin dysfunction"

GLOBAL NETWORK DEGREE (WITHOUT Case4)

| | POST. MEAN | LOWER 95% CI | UPPER 95% CI | EFF.SAMP | pMCMC |
|---|---|---|---|---|---|
| (INTERCEPT) | 1.02438 | 0.36225 | 2.25510 | 3911 | 0.1216 |
| DIAGNOSIS | -1.10485 | 2.16010 | -0.11928 | 8673 | **0.0353*** |
| AGE | 0.01203 | -0.01651 | 0.04085 | 3394 | 0.3880 |

GLOBAL NETWORK STRENGTH (WITHOUT Case4)

| | POST. MEAN | LOWER 95% CI | UPPER 95% CI | EFF.SAMP | pMCMC |
|---|---|---|---|---|---|
| (INTERCEPT) | 1.630103 | 0.361645 | 2.942390 | 8054 | 0.0151 |
| DIAGNOSIS | -0.925860 | -1.880168 | 0.124224 | 9000 | **0.0684** |
| AGE | -0.009228 | -0.034927 | 0.016131 | 9338 | 0.4664 |
