## Supplementary Table4 for "Familial t(1;11) translocation is associated with disruption of white matter structural integrity and oligodendrocyte-myelin dysfunction"

Supplementary Table 4: Comparison of ACTB mRNA levels across lines using RNA-seq and qPCR

| | ACTB RNA-seq (FPKM) | ACTB qPCR (Ct) | GAPDH qPCR (Ct) |
|---|---|---|---|
| Con1 | 863± 71 | 24.81± 0.77 | 24.50± 1.06 |
| Con2 | 984± 240 | 28.38± 1.8 | 25.86± 1.07 |
| Case2 | 1025± 175.2 | 23.12± 0.28 | 24.17± 0.11 |
| Case3 | 697±63.5 | 24.71±1.46 | 25.23± 1.23 |
| Case4 | 745±104.1 | 26.67±1.06 | 26.24± 0.83 |

NB: Measurements were carried out on 3 week oligodendrocyte cultures.
Average FPKM and average Ct values are shown for results derived from RNA-seq qPCR respectively
